# Valence-Based Memory Prioritization in Neural Networks

**DOI:** 10.64898/2026.05.05.723022

**Authors:** Sofiya Zbaranska, Aditya Rajeev, Sheena A. Josselyn, Brokoslaw Laschowski

## Abstract

Improving long-term memory in neural networks remains an open challenge. Existing memory prioritization approaches tend to rely on unsigned importance signals that determine how strongly information should be remembered without representing whether its significance is positive or negative. Inspired by emotional valence in biological memory, we introduce a computational framework that integrates both the magnitude and sign of importance into neural network learning and attention. Our framework combines three mechanisms: (i) a valence-weighted cross-entropy loss that prioritizes salient information during learning; (ii) an amygdala-inspired module that learns structured valence embeddings encoding both magnitude and sign; and (iii) a hippocampus-inspired attention mechanism that incorporates these embeddings into queries and keys to enable valence-sensitive retrieval. We demonstrate generalization of our framework across episodic, spatial, and language-based memory tasks, improving long-term recall of high-salience information. In addition to improving long-term memory, we also show that our framework has the potential to improve memory-based decision-making in reinforcement learning and mitigate the “lost-in-the-middle” problem in language modeling. Taken together, we demonstrate a computational framework that integrates both the magnitude and sign of importance into neural network learning and attention, enabling models to prioritize salient information during learning while using valence-sensitive representations to guide memory retrieval.

## I. Introduction

Despite their success in pattern recognition tasks, artificial neural networks continue to struggle with selective long-term memory storage and retrieval [1], [2]. This limitation is exemplified by the “lost in the middle” phenomenon, where relevant information is disproportionately forgotten as a function of its position in the input sequence [3]. Developing mechanisms that enable machine learning models to selectively prioritize information for long-term memory storage remains an open challenge.

Existing approaches address long-term memory in three main ways. One solution is to extend storage capacity with memory-augmented architectures, as done in Neural Turing Machines, Differentiable Neural Computers, or Transformer-XL models endowed with segment-level recurrence [2], [4], [5]. The second takes inspiration from the brain to change how information is stored in a neural network. For example, [6] showed that nonlinearity derived from N-methyl-D-aspartate receptors incorporated in the feedforward module can improve long-term memory and spatial selectivity in transformers. Principles of Hebbian plasticity have also been integrated into transformers to continuously update and store information through local learning rules [7]. The third approach prioritizes what information is stored using an importance signal. For instance, loss-based reweighting can prioritize experience replay based on temporal-difference error in reinforcement learning [8]. Recently, [9] designed a long-term memory module that preferentially encodes surprising inputs based on gradient magnitude. However, these approaches primarily rely on unsigned importance signals, which determine how strongly information should be prioritized without explicitly representing whether its significance is positive or negative. This limitation motivates a memory mechanism that integrates both the magnitude and sign of importance.

We propose to improve long-term memory prioritization by combining both biologically inspired and prioritization approaches. We introduce a signed importance signal, emulating the role of emotional valence in biological memory. In humans, memory strength follows a V-shaped relationship with the valence of information. Both strongly negative and positive information are recalled more reliably than neutral information (Fig. 1, bottom right) [10]. In the brain, this prioritization is supported by neuromodulators that translate the valence of an event into a chemical signal. Aversive events trigger the release of noradrenaline, whereas rewarding events trigger the release of dopamine, supporting respective emotional associations. The amygdala processes these signals and generates neural activity patterns that encode both the salience of an experience (i.e., its relative importance) and its valence (i.e., whether its emotional affect is positive or negative) [11], [12]. Finally, the hippocampus integrates information across multiple modalities, including the emotional component [12]–[14], which dictates the strength of the formed memory [12]. Biological memory thus relies on a signed signal, in which salience determines how strongly an experience is remembered and the sign of valence helps guide behavior.

**Figure 1.**
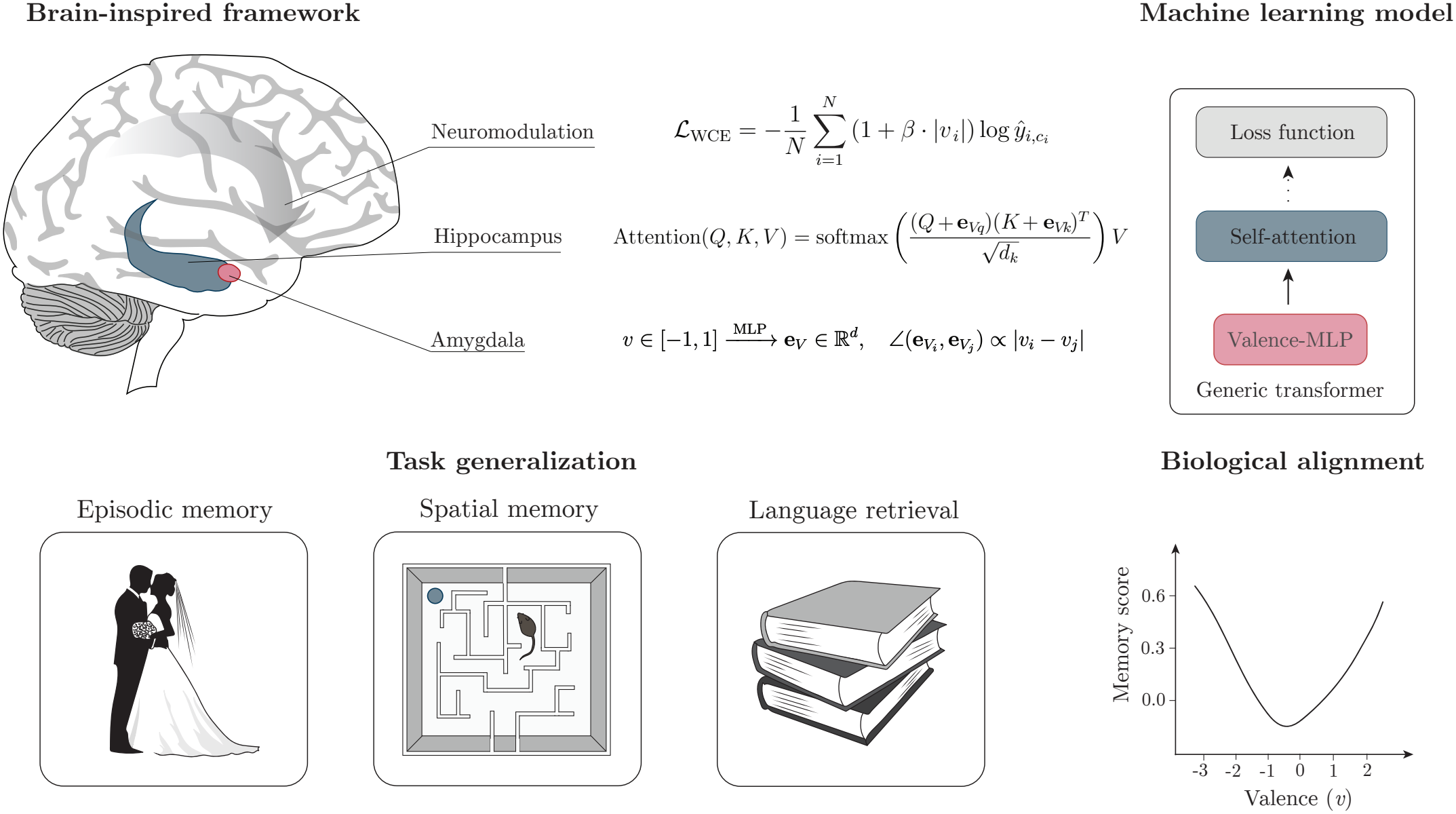
Our computational framework for memory prioritization in artificial neural networks based on emotional valence. Upper row: Neuromodulation is modeled by valence-weighted cross-entropy loss, which scales the learning signal by the valence magnitude (i.e., salience). Hippocampus is modeled as integrating valence embeddings into the attention mechanism to modulate information retrieval. Amygdala is modeled as learning high-dimensional valence embeddings, in which similar valences are aligned and opposite valences are antipodal. Lower row: generalization of our framework across memory tasks (left). Biological memory follows a V-shaped relationship with the valence level (right, adapted from [10]).

**Figure 2.**
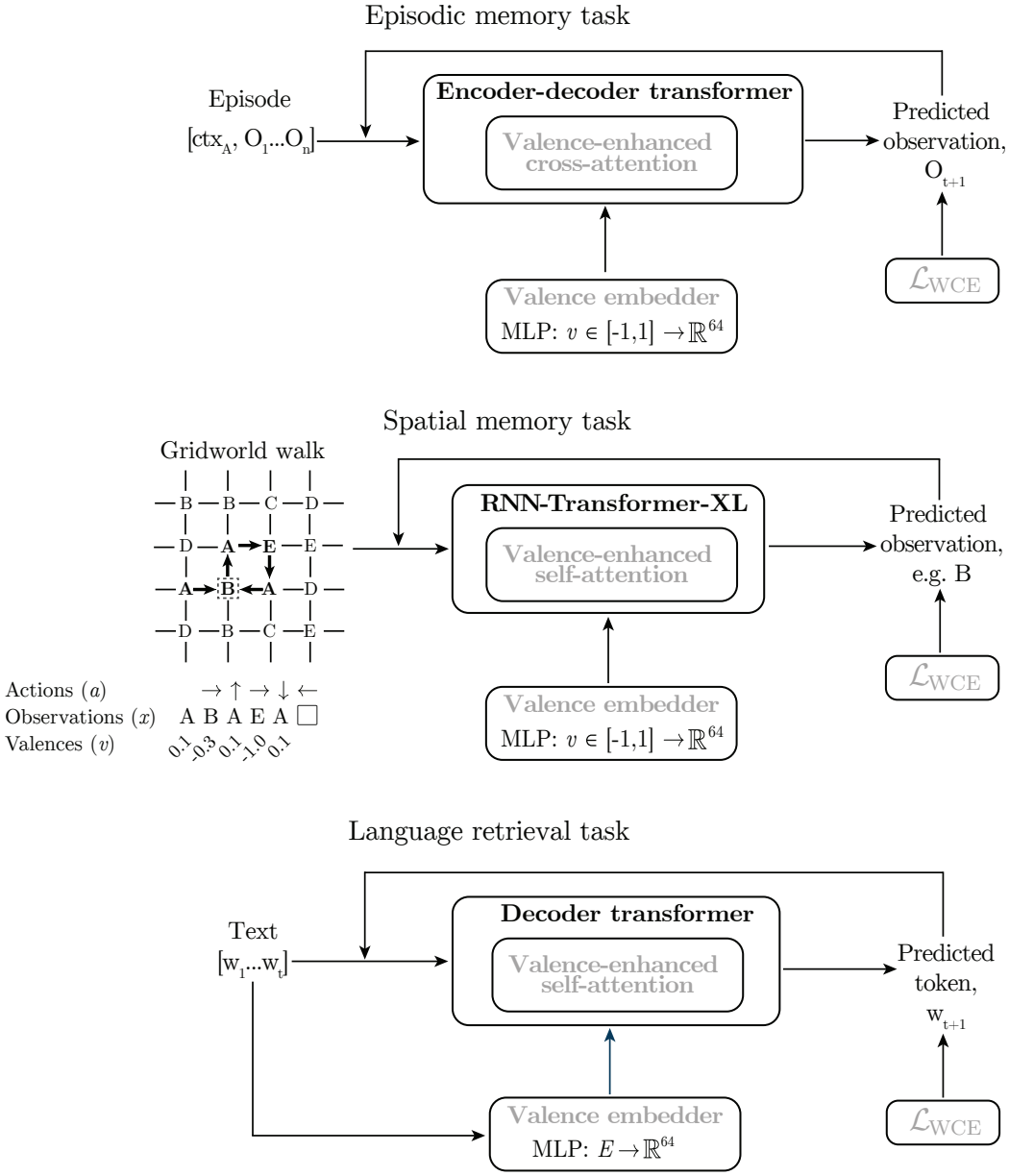
Tasks and models used to evaluate generalization of our framework. Our mechanisms are shown in grey. For the episodic memory task, the model processed sequences of observations paired with a context to predict the next observation given a causal prefix. For the spatial memory task, we trained the model to predict the next observation after taking an action in a gridworld. For the language retrieval task, the model was pre-trained on autoregressive text modeling, then fine-tuned to recall repeated words. Token-level valences were learned from the language model’s token embeddings under NRC VAD lexicon supervision [16] and sentences containing a target word were cued with the model’s own learned valence of that word.

Taking inspiration from these systems-level mechanisms, we developed a computational framework for memory prioritization in artificial neural networks based on emotional valence (Fig. 1). Our framework includes: (i) a valence-weighted cross-entropy loss that scales the learning signal by the valence magnitude ( |*v*|*→* 1: salient, |*v*| *→*0: neutral), analogous to neuromodulation; (ii) an amygdala-inspired mechanism that learns high-dimensional valence embeddings; and (iii) a hippocampus-inspired module that integrates valence embeddings into the attention mechanism to modulate information retrieval. We demonstrate generalization of our framework across episodic, spatial, and language-based memory tasks, improving long-term recall of high-salience information. In addition to improving long-term memory, we also show that our framework has the potential to improve memory-based decision-making in reinforcement learning and mitigate the “lost-in-the-middle” problem in language modeling. Altogether, our contribution is a computational framework that integrates both the magnitude and sign of importance into neural network learning and attention, enabling machine learning models to prioritize salient information while using valence-sensitive representations to guide memory retrieval.

## II. Methods

### A. Framework

Our framework integrates two complementary aspects of valence-based memory prioritization: valence magnitude determines the strength of the learning signal, while both valence magnitude and sign inform memory retrieval through structured embeddings and attention. We implement these functions through three mechanisms that jointly modify learning and memory retrieval.

#### 1) Valence-weighted cross-entropy loss

The first mechanism in our framework is a valence-weighted cross-ntropy loss that scales the learning signal by the valence magnitude, analogous to neuromodulation. We use salience to refer to the valence magnitude |*v*|, which means both strongly positive and strongly negative stimuli are considered salient. In the spatial and episodic memory tasks, ground-truth valence labels *v* ∈ [ −1, 1] were linearly spaced and randomly assigned to each token in a vocabulary, where *v* ≈0 is neutral and *v* = *±*1 maximum salience. This controlled assignment ensures uniform valence frequency across the vocabulary, isolating the effect of valence on memory recall from the confounding influence of observation frequency. We modified the cross-entropy loss to scale each token’s contribution by the absolute valence |*v*_*i*_|:

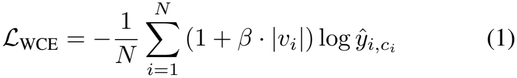

where *β* controls the weighting strength and *c*_*i*_ denotes the ground-truth class.

#### 2) Valence embedder

The second mechanism in our framework is a two-layer perceptron that maps scalar valence *v*_*i*_ to high-dimensional embeddings 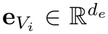, inspired by the amygdala. The embedding norm scales linearly with valence magnitude from 1.0 (neutral) to 2.0 (maximum), while its direction encodes the valence sign. To enforce a geometrically consistent embedding representation, we mapped scalar valence to an angular coordinate 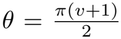, such that *v*=−1 *→* 0,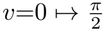 and *v*=+1 *→ π*. We trained the embedder with a geometric contrastive loss *ℒ*_*GC*_ minimizing the mean squared error between the learned cosine similarities and the cosines of the target angular differences:

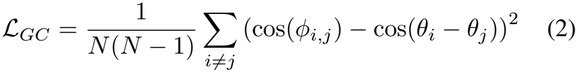

where cos(*ϕ*_*i,j*_) is the cosine similarity between the normalized embeddings **e**_*Vi*_ and **e**_*Vj*_. This encourages embeddings of similar valences to cluster together, while opposite valences (e.g., *v* = 1 and *v* = −1) are pushed toward antipodal directions. We trained the valence embedder in parallel to the main memory stream using the Adam optimizer [15].

#### 3) Valence-enhanced attention

The final module in our framework integrates valence embeddings into the attention mechanism to modulate information retrieval, inspired by the hippocampus. The valence embeddings are projected into the multi-head attention space, mean-centered, and scaled to match the standard deviation of the query and key projections, then added to keys and queries:

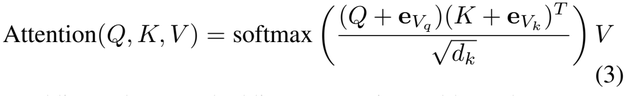

Adding valence embeddings to queries and keys decomposes as follows:

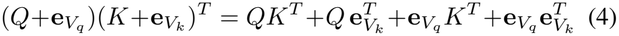

where the first term is standard content-based attention, the cross terms allow content queries to attend to keys of matching valence and vice versa, and the last term captures a signsensitive valence–valence interaction.

### B. Ablations and controls

To evaluate the contribution of each mechanism in our framework, we tested five model configurations: (1) a baseline transformer without valence signaling; (2) weighted cross-entropy (WCE) loss only; (3) WCE with a valence embedder (MLP); (4) the full VALence-Enhanced Transformer (VALET), comprising WCE, valence embedder, and valence-enhanced attention; and (5) a valence embedder with valence-enhanced attention but no WCE. To account for stochasticity, each model configuration was evaluated using three independent seeds. To evaluate generalization, we tested performance across spatial, episodic, and language-based memory tasks. In a reinforcement learning extension of the spatial memory task (Section II-E), we also compared VALET with a salience-only control (SALET) that receives valence magnitude but not its sign.

We further performed two controls to verify whether performance gains arise from the model using valence effectively rather than from its mere presence. First, we trained a baseline model that received valence as an input feature (BL+VF), where the scalar valence of each observation was linearly projected to *d*_model_ and added to its input embedding. Second, we compared our integration of valence embeddings into queries and keys (Eq. 3) with two alternatives. A bilinear bias on the logits (*γ· v*_*i*_*· v*_*j*_, where *γ* is a learned parameter) is a fixed, rank-1 form computed independently of *Q* and *K*. A multiplicative gate *g*(*v*_*i*_, *v*_*j*_) rescales the attention weights according to the sign and magnitude of the query and key valences. These controls were evaluated on the episodic memory task (Section II-C) using ten independent seeds and two-sided Wilcoxon signed-rank tests at each valence level.

### C. Episodic memory task

For the episodic memory task, we created a synthetic dataset mimicking contextual episodic memory, where each episode comprises a context *ctx* ∈ { 1, …, *C }*followed by a sequence of observations *O*_*t*_ ∈ ℝ^256^, each drawn elementwise from *N*(0, 1). We used *C*=32 contexts and *N*_*x*_=1024 observations, with each context assigned a fixed pool of 0.2·*N*_*x*_ observations (sampled without replacement) and associated valences *v*_*t*_∈ [ −1, 1]. One valence level was shared across three different observations. An encoder-decoder transformer [17] served as our baseline with *L*=6 layers in the encoder and decoder, *h*=8 attention heads, *d*_model_=512, and *d*_ff_=2048. The encoder received the full episode *{*`ctx`, *O*_1_, …, *O*_*n*_*}* without masking to form a bidirectional long-term memory representation. The decoder processed the causal prefix {`<SOS>`, *O*_1_, …, *O*_*t*_ *}*autoregressively, with its positional encoding capped at *K*=64 tokens to enforce the working memory horizon. We trained the model to predict *O*_*t*+1_ via cross-entropy loss with teacher forcing using the Adam optimizer (lr = 0.0001) for 100 epochs on 1,000 training episodes (batch 32). Long-term memory retrieval was reported for instances where the target sequence fell outside the *K*=64 step threshold of the decoder’s positional encoding cap, relying on the cross-attention mechanism rather than immediate working memory. Additionally, we quantified the sparsity of the cross-attention weights using the Gini index [18]:

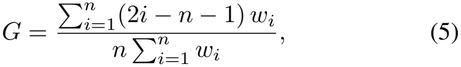

where *w*_1_*≤··· ≤ w*_*n*_ are the attention weights sorted in ascending order, *n* is the number of keys, and *i* is the rank of *w*_*i*_ (*G* = 0: uniform; *G→* 1: maximally sparse). Weights were averaged across heads at each decoding step *t*, and the mean Gini index 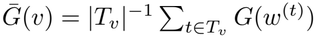 was reported for each valence level *v*, where *T*_*v*_ is the set of steps whose target has valence *v*.

### D. Spatial memory task

For the spatial memory task (adapted from [6]), we modeled the environment as a graph *G*= ( *V,ℰ* ) with |*V*|= *L*^2^ nodes representing discrete locations in an *L× L* grid (*L* = 11). Each node was randomly assigned an observation with a valence *v* ∈ [−1, 1]. The transition probability from node *i* to adjacent node *j* was defined as 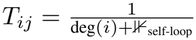, where ⊮_self-loop_ indicates whether a “stay” action was permitted. At each timestep *t*, the agent took an action *a*_*t*_ and had to predict the resulting observation *O*_*t*+1_ from the action and the previously encountered observation-valence pairs {(*O*_*s*_, *v*_*s*_) }_*s≤t*_. Memory retrieval was classified as long-term if the target observation-position pair (*O*_*t*+1_, *p*_*t*+1_) was last encountered more than *K* = 64 steps ago, placing it beyond the model’s context window and thus inaccessible via direct attention over recent tokens. We used an RNN-Transformer-XL [6] as our baseline and trained all models for 200 epochs to predict *O*_*t*+1_ via cross-entropy loss using the Adam optimizer, with the same training parameters as [6]. Similar to Section II-C and previous research [6], we report long-term memory performance for familiar maps.

### E. Reinforcement learning

To test the influence of valence-based memory prioritization on decision-making, we also extended the spatial memory task to reinforcement learning. In each episode, five reward and five hazard locations with valence magnitudes |*v*| ∈ {0.2, 0.4, …, 1.0} were placed randomly in one of the familiar maps from Section II-D. In the exploration trial (750 steps), entering a reward or hazard yielded its value *r*_*t*_ once and each newly visited location yielded a bonus of 0.05. In the recall trial (40 steps), all rewards and hazards were replenished and the agent continued from its last position.

We compared a baseline agent, VALET, and a salienceonly control (SALET) initialized from the same pretrained RNN-Transformer-XL (Section II-D). At each step *t*, a token combining the position code **g**_*t*_, observation, outcome *r*_*t*_, and a trial signal was added to a memory spanning the whole episode. The valence embedder learned valence embeddings 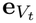 from the experienced value *r*_*t*_ of each location. SALET received only |*r*_*t*|_ such that rewards and hazards of equal magnitude were indistinguishable to attention. Four probe tokens carried the position code of each possible move *a*, obtained by path integration without executing it, attended to the memory without being stored, and predicted the valence *u*_*t,a*_ previously experienced at that location (*u*_*t*,0_ = 0 for staying).

Agents were trained with advantage actor-critic (A2C) [19] and generalized advantage estimation [20] (*γ* = 0.99, *λ*_GAE_ = 0.95, entropy weight 0.01). In each update, 64 episodes were played with the current policy and used to compute the losses.The policy and value were linear readouts of the token’s final hidden state **h**_*t*_:

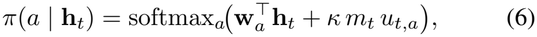

where *κ* is learned and *m*_*t*_ is 0 in the exploration trial and 1 in the recall trial. During exploration, the policy was trained only through A2C on the outcomes and coverage bonus, encouraging exploration. During recall, the probes biased each move towards remembered rewards and away from hazards. The policy was additionally trained towards actions 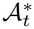 that an exact planner considered optimal given previously experienced locations [21]. The probe and planning losses were

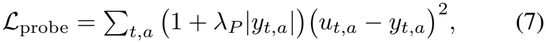

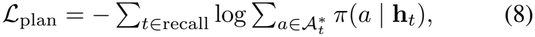

where *y*_*t,a*_ is the valence at the location of move *a* and *λ*_*P*_ = 20 emphasizes remembered rewards and hazards over neutral locations. These were added to the A2C loss, *ℒ*_WCE_ (Eq. 1) and ℒ_*GC*_ (Eq. 2). Each agent was trained for 500 updates with Adam (learning rate 10^−4^ for pretrained and 10^−3^ for new parameters), and evaluated on 512 new layouts identical across agents. We report the net return of the recall trial (cumulative reward plus cumulative penalty) for locations last experienced more than *K* = 64 steps before recall.

### F. Language retrieval task

For the language retrieval task, we used the `Wikipedia (en)` subset of the Pile corpus [22] (100,000 articles with 400 words minimum), split into 70,000 training, 10,000 validation, and 20,000 held-out test articles. Token-level ground-truth valences were derived from the NRC VAD lexicon [16], linearly rescaled to [− 1, 1]. We pre-trained a transformer (*L* = 6, *h* = 8, *d*_model_ = 512, *d*_ff_ = 2048, 512-token context, *∼*50M parameters; BERT-base-uncased tokenization) on autoregressive next-token prediction using AdamW (lr = 10^−4^, weight decay = 0.01, batch size 32), ReduceLROnPlateau scheduling (patience = 3, factor = 0.5), and early stopping (patience = 5). In parallel, the valence embedder learned to map each token’s embedding to a valence embedding via ℒ_GC_ and a scalar regression onto NRC valence, using lexicon words for supervision. We used the same language modeling capability across all models, which were fine-tuned from this shared checkpoint on a repeat-recall task (lr = 3*×*10^−5^, up to 20 epochs, same early stopping). For every lexicon word whose second occurrence was at least *K* = 64 tokens after its first, the loss was computed on predicting the second occurrence (WCE with *β* = 1 where applicable). Every sentence containing a target occurrence was cued with its most salient target word. Models with valence-enhanced attention used the word’s learned scalar valence, where cue strength reflects the model’s own valence estimate. This experiment evaluates valence-guided retrieval of salient words under a targeted cueing protocol, rather than general-purpose language modeling performance. Recall accuracy at the second occurrence was evaluated on the held-out validation set, stratified by the fractional depth of the first occurrence. Similar to previous tasks, Gini index (Eq. 5) was computed from self-attention weights to assess sparsity.

### G. Training and codebase

All models were trained on single NVIDIA H100 GPUs (80 GB GPU memory) with 32 GB of system memory per job, except for our language modeling task, which required 64 GB of system memory. Our codebase for replication is publicly available at https://github.com/neur1s/VALETs.

## III. Results

We investigated whether incorporating both the magnitude and sign of valence into neural network learning and attention improves long-term memory prioritization across different model architectures and tasks. Across episodic, spatial, and language-based memory tasks, our valence-based framework improved long-term recall of highly salient information.

### A. Episodic memory task

#### 1) Memory performance

For the episodic memory task, we trained an encoder-decoder transformer on a synthetic dataset. The model had to recall the correct sequence of observations associated with a context, akin to recalling objects in a room. We evaluated our framework on long-term memory recall and assessed the individual contribution of each mechanism in our general framework (i.e., WCE, valence embedder, and valence-enhanced attention) through systematic ablations.

As shown in Fig. 3A, all models containing WCE displayed a V-shaped relationship between valence and long-term memory accuracy, with curvature depending on the magnitude of *β*. The hyperparameter *β* allows for controllable prioritization: higher values maximize recall of salient observations at the expense of low-salience retention, while lower values yield more modest but balanced improvements. The optimal *β* therefore can be chosen depending on the degree to which high-salience information needs to be prioritized. In our experiment, better recall of high-salience observations (*v* = *±*1) was achieved with WCE alone (for *β≥* 0.5) compared to baseline. However, incorporating all three mechanisms improved recall over WCE alone at all valence levels.

**Figure 3.**
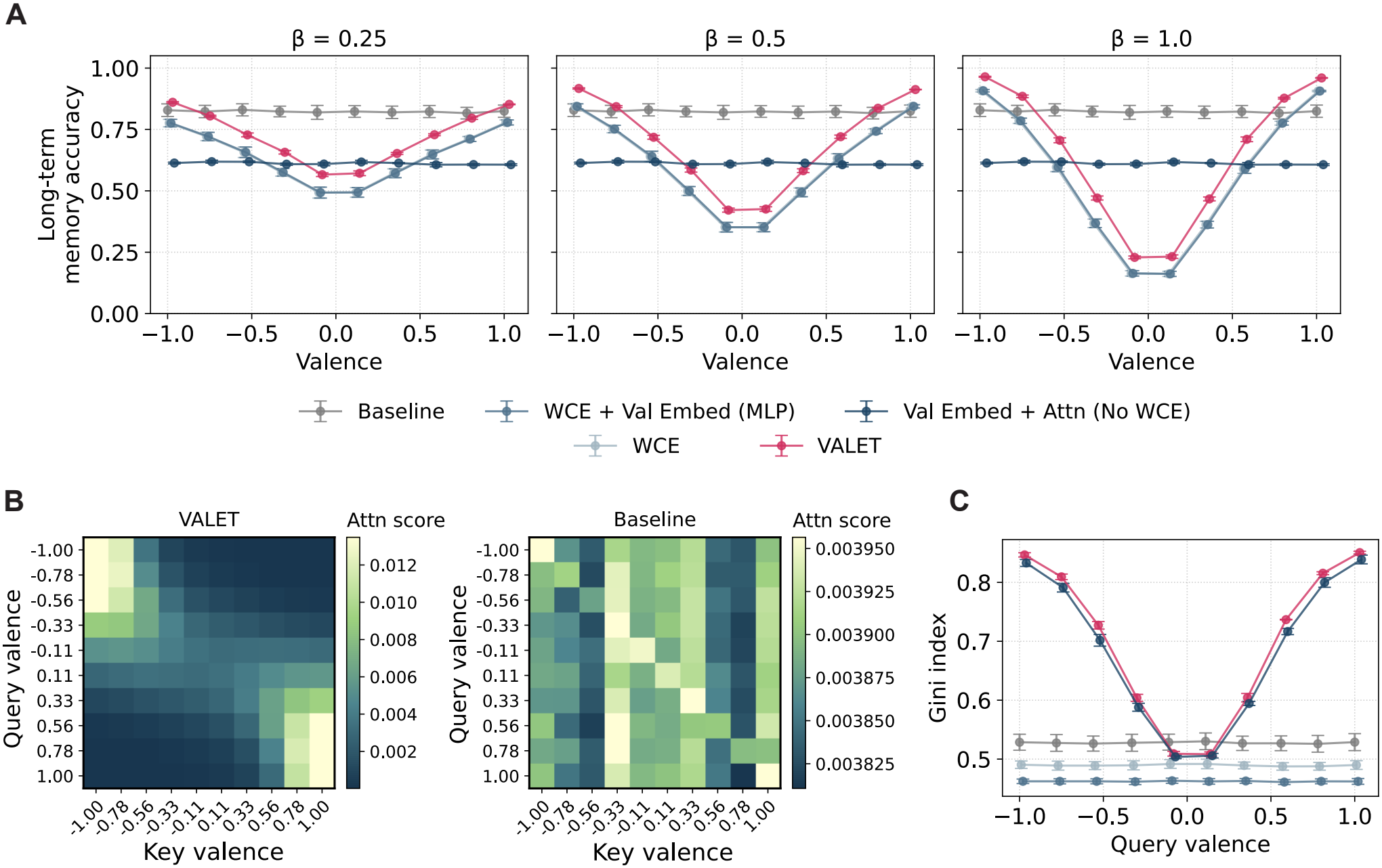
Our valence-enhanced model improved long-term memory recall for high-salience observations on the episodic memory task. (A) Valence vs. long-term memory accuracy across varying levels of *β* (i.e., the strength of learning modulation by valence) for our full valence-enhanced (VALET) model and other ablations. WCE and WCE + Val Embed (MLP) are largely overlapping. (B) Attention score heatmaps grouped by key and query valence levels for our VALET model (*β* = 1) and the baseline. (C) Query valence vs. sparsity of self-attention weights, measured as Gini index (*β* = 1 where relevant). Error bars represent standard error of mean across independently trained models.

Next, we evaluated the effect of our framework on cross-attention scores with respect to observation valence (Fig. 3B). In contrast to the baseline, the cross-attention scores in our full valence-enhanced model were organized by matching valence (i.e., high dot product between maximum valence keys and queries of the same sign) and valence similarity indicated by the gradual fading out along the diagonal (Fig. 3B).

To better understand how our framework improves long-term memory recall of salient observations, we quantified sparsity of the attention weights at the output of the transformer’s cross-attention (Fig. 3C). For VALET (*β* = 1), the Gini index (Eq. 5) exceeded the baseline at all query valences except *v* = *±*0.11, with the gap widening toward high-salience observations (*v* = *±*1). Val Embed + Attn (No WCE) showed a similar V-shaped relationship, indicating that adding valence embeddings to queries and keys modulates retrieval by valence. VALET’s sparsity was marginally higher, suggesting that all mechanisms sharpen attention, likely improving the signal-to-noise ratio and thus recall precision.

#### 2) Design controls

We tested whether VALET’s performance improvement arises from how valence is used rather than from mere access to valence information (Section II-B). We found that providing valence as an input feature (BL+VF) did not change the baseline’s performance at any valence level (Table I). Relative to BL+VF, VALET significantly improved recall of high-salience observations ( |*v*| *≥* 0.78, *p*≤ 0.004). These improvements were accompanied by reduced recall of less salient observations, demonstrating that VALET selectively prioritizes important information rather than uniformly improving memory performance.

**Table I.** Long-term memory accuracy difference (∆) for valence feature control in the episodic memory task. Bl+vf: baseline with valence features, BL: baseline. 10 seeds, Wilcoxon signed-rank test, bold: *p* < 0.05.

|  | BL+VF − BL |  | VALET − BL+VF |  |
|---|---|---|---|---|
| Valence | Δ | *p* | Δ | *p* |
| −1.00 | −0.008 | 0.770 | **+0.199** | **0.002** |
| −0.78 | −0.010 | 0.846 | **+0.124** | **0.004** |
| −0.56 | −0.012 | 0.625 | **−0.051** | **0.027** |
| −0.33 | −0.011 | 0.770 | **−0.296** | **0.002** |
| −0.11 | −0.014 | 0.625 | **−0.530** | **0.002** |
| +0.11 | −0.014 | 0.695 | **−0.532** | **0.002** |
| +0.33 | −0.017 | 0.557 | **−0.291** | **0.002** |
| +0.56 | −0.009 | 0.922 | **−0.046** | **0.037** |
| +0.78 | −0.011 | 0.770 | **+0.120** | **0.004** |
| +1.00 | −0.011 | 0.770 | **+0.199** | **0.002** |

We also compared adding valence embeddings to queries and keys with two alternative forms of valence-enhanced attention: an additive logit bias and a scalar gate (Table II). Both yielded significantly lower recall than VALET for the most salient observations (| *v*| *≥*0.78), with the additive bias also lower at |*v*| = 0.56, whereas differences at lower salience levels were generally smaller and less consistent. Although these differences were small, they were consistent across seeds, indicating that adding valence embeddings to queries and keys prioritizes salient observations more effectively than biasing or gating attention, consistent with the richer content–valence interactions in Eq. 4.

**Table II.** ong-term memory accuracy difference (∆) for alternative valence-enhanced attention mechanisms in the episodic memory task. 10 SEEDS, Wilcoxon signed-rank test, bold: *p* < 0.05.

|  | Additive bias − VALET |  | Scalar gate − VALET |  |
|---|---|---|---|---|
| Valence | Δ | *p* | Δ | *p* |
| −1.00 | **−0.007** | **0.002** | **−0.010** | **0.002** |
| −0.78 | **−0.015** | **0.002** | **−0.018** | **0.002** |
| −0.56 | **−0.015** | **0.002** | −0.009 | 0.193 |
| −0.33 | −0.006 | 0.232 | +0.010 | 0.322 |
| −0.11 | −0.004 | 0.164 | +0.009 | 0.275 |
| +0.11 | −0.005 | 0.232 | +0.010 | 0.232 |
| +0.33 | **−0.006** | **0.037** | +0.008 | 0.432 |
| +0.56 | **−0.018** | **0.006** | −0.013 | 0.232 |
| +0.78 | **−0.012** | **0.002** | **−0.015** | **0.002** |
| +1.00 | **−0.006** | **0.004** | **−0.009** | **0.002** |

### B. Spatial memory task

To further evaluate the generalization of our framework on long-term memory recall, we trained an RNN-Transformer-XL model [6] on a gridworld memory task in which the model had to correctly recall observations at locations visited outside the context window. Long-term memory accuracy varied depending on the model and hyperparameter *β* (Fig. 4A). All models containing WCE showed a V-shaped relationship between valence and memory accuracy, with the gradient of the curve depending on the size of *β*, which specifies the strength of loss weighting by |*v*| (Fig. 4A). Including just one module (e.g., either WCE or valence-enhanced attention) was insufficient to improve recall of high-salience observations compared to baseline. This required the combination of all three mechanisms in our framework, as shown in our VALET model.

**Figure 4.**
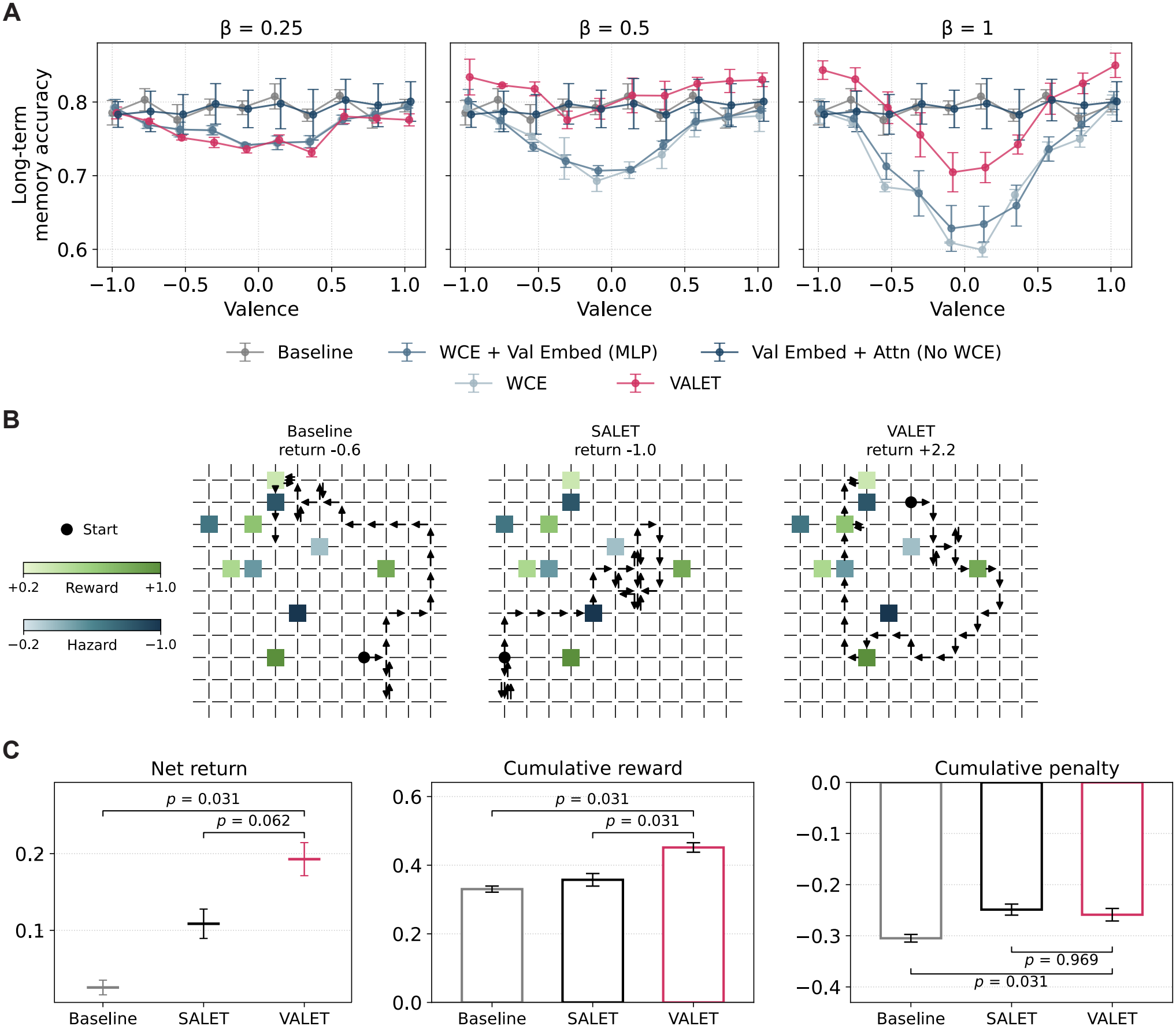
Our valence-enhanced model improved long-term memory for salient observations and reinforcement learning-based decision-making in the spatial memory task. (A) Long-term memory accuracy as a function of valence across varying levels of *β* modulation strength for our VALET model and other ablations. (B) Example recall trajectories of the baseline, salience-only (SALET), and VALET agents on the same map after exploration. Reward (green) and hazard (blue) locations are shaded by valence magnitude and the filled circle represents the start of the recall trial. (C) Net return and cumulative reward and penalty in the recall trial for rewards and hazards last experienced more than 64 steps earlier (long-term memory). Error bars represent standard error of mean across independently trained models and *p*-values are shown for one-sided Wilcoxon signed-rank tests (*n* = 5 seeds).

### C. Reinforcement learning

To test the utility of valence-based memory prioritization in cases where discriminating the sign may be important for task performance, we extended the spatial memory task to decision-making with rewards and penalties using reinforcement learning (Fig. 4B). First, the agents explored maps with five rewards and hazards, then had to maximize their return by collecting rewards and avoiding hazards from memory (Section II-E). VALET achieved the highest net return, exceeding the baseline and showing a tendency toward a higher net return than the sign-agnostic SALET (Fig. 4C). Decomposition of the net return revealed a dissociation between salience and valence. VALET obtained a higher cumulative reward than SALET and baseline. However, the cumulative penalty was comparably low in SALET and VALET. Both incurred less penalty than the baseline, indicating that hazard avoidance was similarly effective with or without the sign of valence. These findings suggest that salience-based memory prioritization supports hazard avoidance, whereas preserving valence sign can provide additional benefits for retrieving rewarding locations and guiding memory-based decisions.

### D. Language retrieval task

To evaluate the generalization of our framework beyond synthetic memory tasks, we investigated whether valence-enhanced learning and attention improve long-range retrieval in language-based memory tasks. We developed a task in which a pre-trained model had to recall a previously encountered word at its second occurrence. Valence was learned from token embeddings rather than directly supplied. The valence embedder was trained under supervision from human-annotated valence ratings in the NRC VAD lexicon [16]. The model learned a geometrically consistent valence representation, where cosine similarity between embeddings decreased gradually with increasing difference in NRC valence (Fig. 5A).

**Figure 5.**
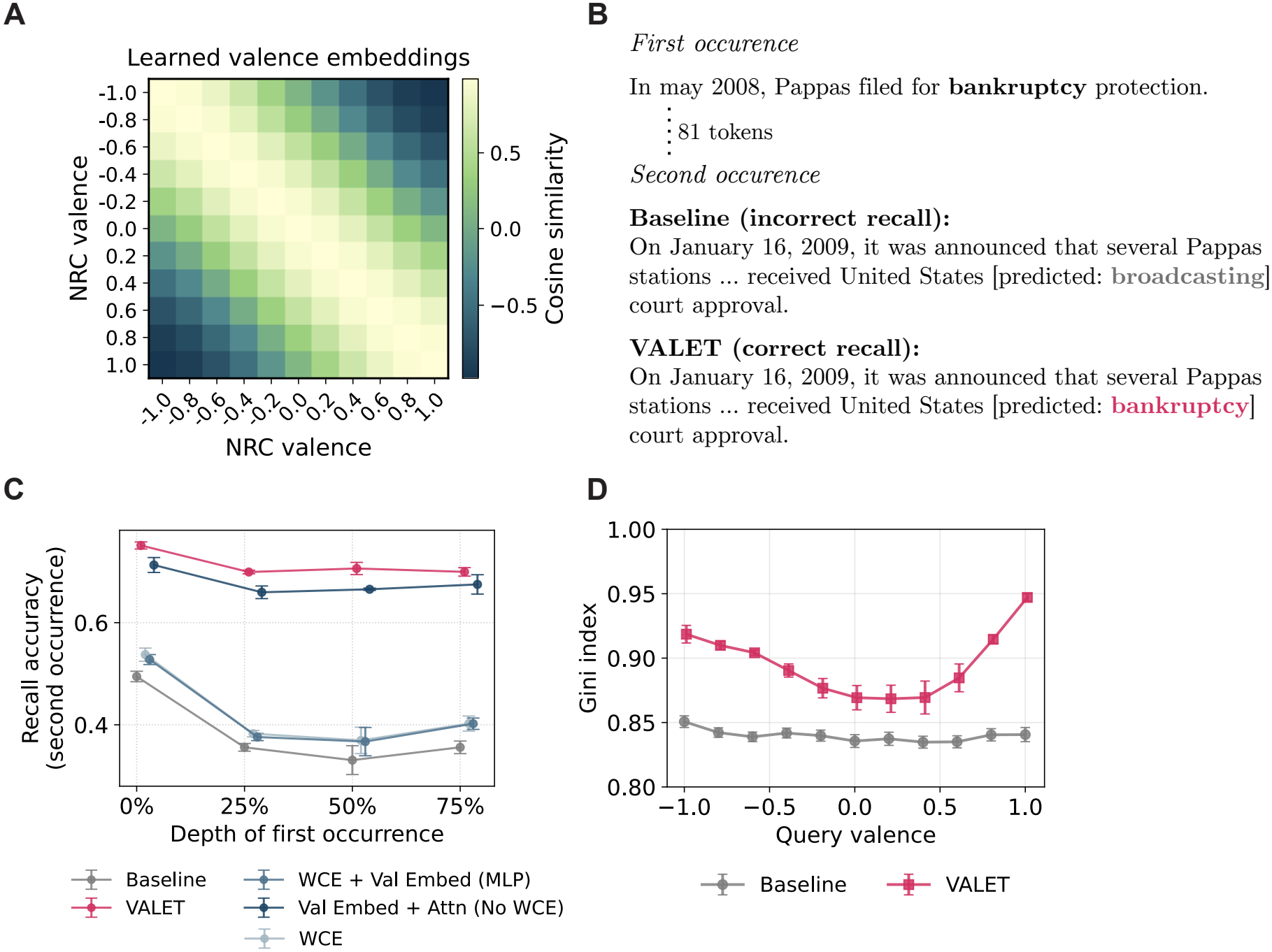
Our valence-enhanced model achieved improved recall of high-salience words across context depths in language modeling. (A) Cosine similarity heatmap of valence embeddings learned from token embeddings. (B) Example showing the baseline failing to recall the previously encountered word “bankruptcy” (NRC valence -0.98), while our VALET model reliably recovered the correct word. (C) First-occurrence depth vs. recall accuracy for VALET and other ablations for words with |*v*| *≥*0.7. (D) Attention weight sparsity (Gini index) increases toward extreme valence levels in the VALET model, but not the baseline. Error bars represent standard error of mean across independently trained models.

We evaluated recall of high-salience words ( |*v*|*≥* 0.7) as a function of the depth of their first occurrence (0%, 25%, 50%, and 75%), with the query positioned within the last 10% of the context to emulate the “lost-in-the-middle” problem in language modeling. The baseline showed a characteristic decline, dropping from 0.49 at 0% to 0.33 at 50% with a partial recovery at 75% (Fig. 5C). VALET (trained with *β* = 1, based on previous results) outperformed the baseline at every depth by nearly 0.3 and flattened the depth-accuracy curve. Ablations showed that WCE improved recall only modestly, whereas valence-enhanced attention contributed to the largest improvement. The full VALET model performed best at all depths. Qualitatively, the baseline often predicted semantically plausible but incorrect words at the second occurrence, while VALET recovered the correct high-salience word (Fig. 5B). At recall, VALET concentrated its attention more sharply than the baseline, with the Gini index of the attention weights following a V-shaped relationship that peaked for both strongly negative and positive queries (Fig. 5D). Overall, these findings demonstrate that valence-enhanced attention improves retrieval of salient tokens under the targeted cueing protocol in our experiment.

## IV. Discussion

In this study, we developed a computational framework for memory prioritization in artificial neural networks based on the principle of emotional valence. We introduced a signed valence whose magnitude reflects salience and whose sign distinguishes positive from negative affect. This signal informs three mechanisms: valence-weighted cross-entropy loss, a valence embedder, and valence-enhanced attention. Our approach differs from previous studies that have focused on synaptic-level plasticity [6], [7] or unsigned importance signals to improve long-term memory in machine learning [8], [9]. Across different memory tasks and model architectures, our framework improved longterm recall of both strongly negative and positive information. More broadly, our results suggest that valence-based prioritization can influence both which information neural networks retain and how they retrieve it.

Our findings extend previous approaches in three important ways. First, our valence signal produced a V-shaped relationship between valence and memory recall, characteristic of biological memory [10], rather than uniform improvement in [8] and [9]. Our ablations suggest that prioritization benefits from combining valence-weighted cross-entropy loss and valence-enhanced attention, with their relative contributions varying across tasks. Second, our valence embedder learns embeddings that are added to queries and keys before attention is computed. Previous research has modified attention by adding fixed biases to the attention logits [23]. Our approach achieved higher recall of salient information than both an additive logit bias and a multiplicative gate (Table II). Third, although memory performance was similar for both strongly negative and positive stimuli, attention was organized by valence similarity, with the highest scores between salient queries and keys of the same sign and the lowest between opposite signs (Fig. 3B).

Our reinforcement learning extension further suggested that this distinction can benefit decision-making: VALET collected more rewards than the salience-only model, although both avoided hazards similarly (Fig. 4C). This distinction is consistent with findings that the amygdala contains distinct neuronal populations associated with positive and negative valence that influence corresponding behaviors [24].

Beyond synthetic memory tasks, we showed that our valence-based prioritization framework has the potential to mitigate the “lost-in-the-middle” problem in language modeling, characterized by decreased retrieval accuracy for information in the middle of long sequences [3]. While previous studies have relied on extensive fine-tuning [25] or external key-value stores [26], our framework modulates memory retrieval through valenceenhanced attention. Behrouz et al. [9] similarly used an importance signal based on surprise to improve language modeling, although their focus on next-token prediction precludes direct comparisons. Our valence embedder learned structured valence embeddings from token embeddings under supervision from the NRC VAD lexicon [16]. Using a 50M-parameter pretrained language model fine-tuned for repeated-word retrieval, our VALET model improved recall of salient words across all tested context depths, flattening the depth-accuracy curve.

Despite these developments, our research serves as a proof-of-concept and is not without limitations. We used synthetic episodic and spatial memory tasks with valence assigned uniformly at random to isolate the effect of valence on memory from confounds such as observation frequency or semantic similarity. This design does not reflect the valence distributions of naturalistic datasets. Only the language task used valences learned from real-world text under supervision from human-annotated valence ratings. Our framework is also mainly applicable to settings with meaningful valence structure, such as affective contexts, rewards, or action outcomes, and may not benefit purely neutral settings. We showed that stronger prioritization at high *β* reduced recall of neutral information (*v≈* 0). This trade-off must be considered according to the semantics of the task, depending on how strongly salient information needs to be prioritized over neutral information. In the language retrieval task, recall was cued with the model’s learned valence, which was available only to models with valence-enhanced attention. Therefore, the improvements reflect targeted valence-guided retrieval rather than general-purpose language modeling performance. Finally, our context length of 512 tokens is shorter than the contexts of 2,000–16,000 tokens in which the lost-in-the-middle effect was originally reported [3] and was chosen to match the model size used in our other tasks.

These limitations motivate future research. Beyond synthetic memory tasks and language modeling, valence-based prioritization may be particularly useful for embodied agents that process continuous visual streams and must choose actions [27]. Outcomes such as reaching a goal or colliding with an obstacle are highly positive or negative, and both are important to remember. Assigning valence to observations based on the outcomes of subsequent actions could allow an agent to prioritize the frames that matter most for later decisions. Another important direction is to scale our framework to pre-trained large language models, for example by training the valence embedder and low-rank adapters on the query and key projections while keeping the rest of the model frozen. Valence could be learned not only through lexicon supervision but also from reward signals [28] or human preference ratings. Large language models may already encode valence implicitly, since sentiment has been shown to be linearly represented in their activations [29]. Our valence embedder could learn valence from token embeddings alone, indicating that valence structure is already present in the language data. Future studies should investigate whether making valence explicit through valenceweighted cross-entropy loss and valence-enhanced attention improves the ability of large language models to retrieve salient information. Finally, the system-level grounding of our framework leaves open the question of mechanistic alignment [30]. It remains unknown to what extent the internal representations in our valence-enhanced models align with patterns of activity in biological neural networks during emotional memory encoding and retrieval.

## V. Conclusion

In conclusion, we developed a computational framework for memory prioritization in artificial neural networks that integrates both the magnitude and sign of valence into learning and attention. Our framework combines valence-weighted cross-entropy loss, structured valence embeddings, and valenceenhanced attention to prioritize salient information during learning and guide memory retrieval. Across episodic, spatial, and language-based memory tasks, our model improved long-term recall of highly salient information, with ablation experiments demonstrating the benefits of combining these mechanisms. We further showed that our framework can improve memory-based decision-making in reinforcement learning and mitigate the “lost-in-the-middle” problem in language modeling. Overall, these findings highlight the potential of valence-based memory prioritization to improve selective long-term memory retrieval across different neural network architectures and tasks.

## VI. Acknowledgements

We thank members of the Machine Intelligence Lab for their insightful discussions and feedback. We dedicate this research to students and researchers in Ukraine. Their resilience and unwavering commitment to education and learning continue to serve as a beacon of hope and inspiration to the global academic community.

